# ‘The Bold are the Sociable’: Personality, sociability and lateralized utilization of brain hemisphere in the juveniles of a megafish Deccan Mahseer (*Tor khudree*)

**DOI:** 10.1101/683532

**Authors:** Vishwanath Varma, Harsh Vasoya, Anushka Jain, VV Binoy

## Abstract

The present study explored relationships between personality traits; boldness, activity and sociability, and lateralized utilization of brain hemispheres in the hatchery reared juveniles of Deccan Mahseer (*Tor khudree*), a game fish inhabiting the rivers of central and southern India. Our results revealed a significant positive correlation between boldness and activity in this species when tested in isolation. However, boldness was positively correlated with the time spent near the individual conspecific but not with the individual alien invasive heterospecific tilapia (*Oreochromis mossambicus*). Although juvenile Deccan mahseer exhibited significant variation in the preference towards conspecific over heterospecific, no divergence in the utilization of right or left eye was seen while observing these individuals suggesting the lack of lateralized utilization of the brain hemispheres. Furthermore, laterality in visual preference failed to show any significant correlation with any of the personality traits tested in this species. Results are discussed in the light of the existing literature on the impact of life in homogenous hatchery conditions on the behaviour, personality traits and cognitive abilities of fishes.

**Significance statement:** The present study is one of the first that focuses on personality and lateralization in Deccan Mahseers, an endangered freshwater megafish. We report a positive linkage between boldness and sociability but do not find any correlation of personality with lateralized utilization of brain hemispheres in diverse social contexts. These findings have implications in the conservation and cultivation of this ecologically, culturally and economically important indigenous fish.

## Introduction

Inter-individual variability despite experiencing similar environmental conditions is commonly seen in many phenotypic traits within several animal species. Some behavioural traits exhibit consistency across time and contexts and constitute the personality of an individual (Reale et al. 2007). Behavioural consistency and the tight linkage between various behavioural (behavioural syndrome) and physiological traits can limit behavioural flexibility - the capacity to modulate behaviours in accordance with the demands of the situation - at the level of the individual or population (Biro and Adriaenssens, 2013; Villegas-Rios et al., 2018) and affect their fitness (Smith and Blumstein, 2008; Le Galliard et al., 2013). The determinants of origin, maintenance and modification of personality traits such as boldness, exploration, aggression, activity and sociability (also known as big five animal personality traits, Conrad et al. 2011) have been explored across various taxa of fishes, one of the most widely studied animal groups for such variation in the behaviour. The distribution of these traits in populations of piscine species inhabiting diverse natural and artificial habitats, syndromes between these traits (Sih et al. 2004) and its impact on various cognitive abilities such as learning and memory, ability to discriminate between shoals of different sizes, use of social information, etc. (Carere and Locurto, 2011; Trompf and Brown, 2014; Lucon-Xiccato and Dadda, 2017) have also attracted the attention of researchers across the world.

Although individual variation and its consistency are being studied at behavioural and physiological (e.g. metabolic rate) levels (Boulton et al., 2015; Rupia et al., 2016; Mathot and Frankenhuis, 2018), search for its neural correlates has received greater attention in the last two decades (Coppens et al., 2010; Silva et al., 2014; Soares et al., 2018). According to Goursot et al. (2018), studying the relationship between personality traits and the lateralized utilization of brain hemisphere could provide insights into the neurobiological mechanisms underlying individual variation in behaviour. Many species of vertebrates and invertebrates exhibit a preference to use one cerebral hemisphere over the other while performing certain cognitive tasks and such lateralization can also be observed in choosing left and right locomotor organs while performing an activity (Frasnelli and Vallortigara, 2018). Due to their morphological and neuroanatomical uniqueness resulting in the laterally positioned eyes generating largely independent images due to minimal overlap in the visual fields of the two eyes and each eye sending information to the contralateral brain hemisphere, fishes have been favourite model systems of lateralisation researchers (Larsson, 2015; Hori et al. 2017). Interestingly, similar to personality, laterality is also not consistent across populations (Brown et al., 2004) and inter individual variation in this trait has been reported in many piscine species. Some individuals prefer to use left eye (hence right hemisphere) to observe a stimulus, while others may use right eye (Bisazza et al., 1997; Brown and Bibost, 2014). Individuals without any strong preference for eye use are also common in fish populations (Brown et al., 2004; 2007; Takeuchi and Hori, 2008). According to Frasnelli and Vallortigara (2018), individuals vary not only in the bias for utilising brain and locomotor organs but the strength of lateralization as well, and such divergence may be related to their various cognitive abilities.

The relationship between laterality and personality in fish continues to be a hot topic of debate and some researchers consider personality as a determinant of biased usage of brain hemispheres while others believe these two vital traits may have coevolved (Reddon and Hurd, 2009; Irving and Brown, 2013). Empirical studies using various fish revealed that relationship between personality and laterality is sensitive to the species, sex, life experiences of the individual, etc. (Irving and Brown, 2013; Brown and Bibost, 2014). For instance, bold convict cichlids (*Amatitlanianigrofasciata*) exhibited stronger lateralisation (Reddon and Hurd, 2009); whereas these traits were not found to be correlated in Port Jackson shark (*Heterodontus portusjacksoni;* Byrnes et al. 2016) and guppies (*Poecilia reticulata;* Irving and Brown 2013). However, studies showing the positive correlation between the strength of lateralisation with emotional and stress response in various animals including fishes (Goursot et al. 2018; Byrnes et al. 2016) points to the need for extending this line of research to more piscine species especially those experiencing changes in the quality of their habitat. The knowledge of individual and population variation in the lateralization and its connection with vital behaviours such as boldness, activity and sociability and their impact on behavioural adaptations to the social and ecological changes experienced by the individual would be useful in understanding the mechanism of action of the evolutionary selection pressures on these vital behavioural and neurobiological traits. Additionally, such information could also contribute significantly to devise population specific strategies to ensure the wellbeing of the individuals living in captive conditions or released to the natural habitats as part of the reintroduction or restocking programmes (Huntingford and Adams, 2005; Galhardo and Oliveira, 2009; Conrad et al., 2011; Castanheira et al., 2017).

In India, populations of Deccan Mahseer (*Tor khudree*), a mega fish found in the natural water bodies of the northern Western Ghats, growing up to 40 kg in size (Nautiyal, 2014), are in steady decline due to high anthropogenic pressure and habitat loss (Kharat et al. 2003, Raghavan et al. *in press*). However, various governmental and non-governmental agencies are working in this area with an aim to protect the existing mahseer populations. The hatchery run by the Department of Fisheries, Government of Karnataka, India located at Madikeri, Coorg, breeds this fish in captivity and the young ones are released into the rivers and dams every year. Although not studied in the Deccan mahseer, research has shown that hatchery reared fishes struggle to survive in the natural habitats post release due to behavioural deficits (Brown and Laland, 2001). Moreover, living in homogenous environmental conditions lacking ecological selection pressure, such as the rearing tanks in hatcheries, could alter personality traits and lateralisation in fishes (Brown and Day, 2002; Sundstrom et al., 2004; Bibost et al., 2013; Sykes et al., 2018; James et al., 2018) and hence could negatively impact the success of restocking interventions. The situation is further complicated in the case of juvenile Deccan mahseers released into the natural water bodies where they have to face growing populations of Alien Invasive Piscine Species (AIPS) such as Tilapia (*Oreochromis*), grass carp (*Ctenopharyngodon idella*), common carp (*Cyprinus carpio*), etc. According to Blanchet et al. (2007) interaction with the alien invasive species could modify several vital behaviours of indigenous fishes which could alter selective pressures and move them towards extermination. Although provision of behaviour based life skill training to the juveniles could increase their capacity to adapt to the novel environment to which they are released (Brown and Laland, 2001; Hyvarinen and Rodewald, 2013; Sloychuk et al., 2016), unfortunately, no information on the behaviour and personality traits of the individuals of Deccan mahseers living in captivity or natural water bodies or the nature of their interaction with indigenous or alien invasive heterospecifics is available in the literature.

The present study explored the following questions:

a. Is there any linkage between three major personality traits, boldness, sociability and activity of the hatchery reared juveniles of Deccan mahseer?
b. Does this species show the syndrome between personality and lateralized utilization of brain hemisphere in various social contexts?
c. Do they express similar levels of sociability and laterality towards a conspecific and an individual of AIPS tilapia?

## Methods

### Test animals and husbandry

Fingerlings of Deccan Mahseers (4 ± 1.3 cm Standard Length; SL ± SD and 2 months old) were procured from the Karnataka Fisheries Department hatchery, Kodagu, Karnataka State, India. The heterospecific, Mozambique tilapia (*Oreochromis mossambicus*) were purchased from a professional aquarium keeper in Bangalore and were of similar size to the subject mahseers. In the laboratory, groups of 8 individuals were made in aquaria of size 40 •25 • 27 cm. Three sides of these tanks were covered with black paper to visually isolate the groups from each other. Steel grills wrapped with mosquito nets were placed over the tanks to prevent fishes from jumping out. The water level was kept at a height of 20 cm, temperature was maintained at 25 ± 1 °C along with a light regime of 12:12 hour (L:D light-dark cycle). Fishes were fed *ad libitum* with commercial food pellets (Taiyo fish food) every day in the evening. The tanks were aerated throughout the day to maintain good oxygen level and the water was changed after every 10 days. The Tilapia were also maintained and fed in a similar manner.

### Testing personality traits

Personality traits of the subject fishes were quantified using swim-way apparatus (Yoshida et al., 2005; Binoy, 2015). This instrument comprised of an aquarium (80 • 40 • 33 cm) divided into two unequal chambers using an opaque black Plexiglass sheet; chamber A (15 • 40 • 33 cm) the small compartment was used as start chamber, and chamber B (65 • 40 • 33 cm) was the swim-way for exploration. The partition wall was provided with a gate (8 • 5 cm) which could be closed using a guillotine door. The apparatus was covered with black paper on all four sides and the floor was white coloured to provide greater visibility of the subject fish in the video of the experiment recorded for the analysis. The water level was maintained at 15 cm and a compact fluorescent lamp (40 W) hanging on top illuminated the experimental arena. The apparatus was kept inside a section of the room cordoned off by black curtains to avoid external disturbances and a video camera fitted above was used for recording the experiments.

The subject fish was introduced into chamber A and the guillotine door was raised after 5 minutes given for acclimation. Shy fish are known to spend more time in the start chamber in comparison to their bold counterparts, as it could function as a shelter as opposed to the open and well-lighted area of the swim-way (Toms et al., 2010). The individual fish was given 5 minutes for the exploration of chamber B and ‘emergence latency’ as a measure of their boldness was recorded. Those individuals that remained in the start-chamber after this period were given ceiling values of ‘300 seconds’ and the trial was terminated. Sixty two individual fishes were tested following this protocol and time taken by the fish to come out of the start chamber (boldness), time spent actively moving in the swim-way and the frequency of switching chambers and crossing the central area of the swim-way were quantified from the video record using BORIS v.6.2.2 (Behavioral Observation Research Interactive Software; Friard and Gamba, 2016).

### Testing sociability and lateralized utilization of the brain hemispheres

The apparatus for studying sociability and lateralization of individual mahseer was an aquarium (80 • 40 • 33 cm) divided into two compartments (chamber A and B respectively) connected by a narrow channel of the dimension 44 • 15 • 33 cm (Bisazza et al., 1997). Here also compartment A (14 • 40 • 33 cm) was used as the start chamber and the entry of the subject fish from it into the narrow channel was controlled using a guillotine door. Chamber B functioned as the detour chamber (22 • 40 • 33 cm) and a transparent stimulus cage (13 • 10 • 33 cm) kept in it was used to present the stimulus fish. All partition walls of this apparatus were made using opaque Plexiglass, and similar to the open field apparatus, this apparatus was also isolated from external disturbances by covering all sides using black paper. Here also water level was maintained at a height of 15 cm.

In order to test the lateralized utilization of brain hemispheres and sociability, the subject fish which had undergone personality test were introduced into the start chamber of the apparatus individually. After 5 minutes of acclimatization, the guillotine door present between chamber A and the narrow channel leading to the chamber B was raised. Individuals that did not enter the narrow channel were driven inside using a hand net (Bisazza et al., 1997). After reaching the chamber B, the fish was given 5 minutes to explore the stimulus in the presentation cage. Each subject was separately tested for the lateralized utilization of brain hemisphere and sociability towards three different cues, unfamiliar conspecific, unfamiliar heterospecific (tilapia) and empty stimulus cage (control). After each trial, the subject fish was moved from the experimental arena to another tank and a rest period of 10 minutes was provided. The sequence of the presentation of the stimuli across subject fish was randomized. Water in the apparatus and the conspecifics used as stimulus fish were changed after each trial while the heterospecifics were repeated after every 5 trials. All fishes that were tested for personality (n = 62), except one individual which did not enter the detour chamber, were used for assaying sociability and lateralized utilization of brain hemisphere as well. Sociability was measured as the time spent by the focal fish within 2 cm of the stimulus cage and laterality, as the duration of using left and right eyes for observing the stimulus given during the test period. If the head of the subject fish was perpendicular to the partition or when it formed an angle greater than 90° with respect to the partition separating the stimulus chamber, the fish was not considered to be viewing the stimulus with left or right eye (Sovrano, 2004). Laterality Index (LI) was calculated using the formula LI = ((Duration using Left eye-Duration using Right eye)/(Duration using Left eye+Duration using Right)) ×100 as described in Bisazza et al. (1997).

### Analysis

The three response variables of activity, i.e. total activity, number of times crossing centre and number of times switching between chambers were first analyzed separately with a ‘full’ model (Generalized Linear Models) using other personality traits as covariates. Subsequently, non-significant main effects were deleted stepwise to obtain the most parsimonious model with the best fit (Crawley, 2007). When total activity was used as response variable, latency to emerge into novel environment (boldness) and number of times crossing the centre were taken as covariates. For the other two response variables (i.e. number of times crossing centre and the frequency of switching chambers), boldness and activity (total time spent swimming in open field) were taken as covariates. Lateralization and sociability were also used as response variables for mixed models where boldness and activity were used as covariates. The type of stimulus (controls, conspecific and heterospecific) was used as fixed factor while FishID (which indicates the identity of individual fish) was taken as random factor for analysis using linear mixed models (LMM). Models were compared using the Akaike Information Criterion where models with lower AIC values are considered better fits of the dataset. Coefficients of slope of individual effects of factors and interactions are depicted by β and mentioned when the slopes were statistically significant at p < 0.05. Models were also checked for homogeneity of variance by visual inspection of the residual plots (Kelley et al., 2008; Maskrey et al., 2018). Total activity and sociability data were log-transformed since they showed heteroscedasticity with an increase in variance along with the mean as determined from the residual plots. Frequency of switching chambers was subjected to GLM with the poisson family and log link function since they represent count data. However, since number of times crossing centre showed overdispersion in the residuals, GLM of quasipoisson family was chosen for this response variable.

One-sample Wilcoxon signed-rank test was performed on laterality index scores against zero values to determine the intrapopulation variation since the data did not follow normal distribution after transformation (Kolmogorov–Smirnov test). Kruskal-Wallis test was performed to compare the sociability and laterality in order to test cross-context variation (empty cages, conspecifics and heterospecifics) in these traits. Non-parametric Spearman’s rank correlation was used for testing the relationship between various parameters measured.

All statistical tests were performed on R ver. 3.5.1 using ‘glm’, ‘lme4’ and ‘lmerTest’ packages while Spearman’s rank correlation shown in figure legends was conducted on STATISTICA version 7.0.

## Results

### Personality traits of Deccan Mahseer

Full model of total activity (log transformed) as response variable with boldness and number of times crossing the centre as covariates were supported (as evidenced by lower AIC values) over models involving only number of times switching between the shelter (ΔAIC = 70.8) or only boldness (ΔAIC = 1.5) as covariate. The full model yielded a negative estimated coefficient of latency to emerge on total activity (β = −0.007; p < 0.001) suggesting that bold individuals show greater activity (Figure 1a). However, no significant effect of crossing centre on activity (β = 0.022; p > 0.05) was observed. Modeling of the frequency of crossing the centre as response variable (with quasipoisson distribution) showed statistically significant effect of boldness (β = −0.006; p < 0.05; Figure 1b) as covariate but no significant effect of total activity (β = 0.004; p > 0.05). However, switching the chambers (with poisson distribution) appeared to be significantly correlated with both boldness (β = −0.019; p < 0.001; Figure 2a) and activity (β = −0.007; p < 0.001; Figure 2b) since the full model revealed lowest AIC compared to models involving only either of the two as covariates. All models were better fits than their corresponding null models. Hence, multiple components of personality of the Mahseers were correlated with one another.

**Figure 1.**
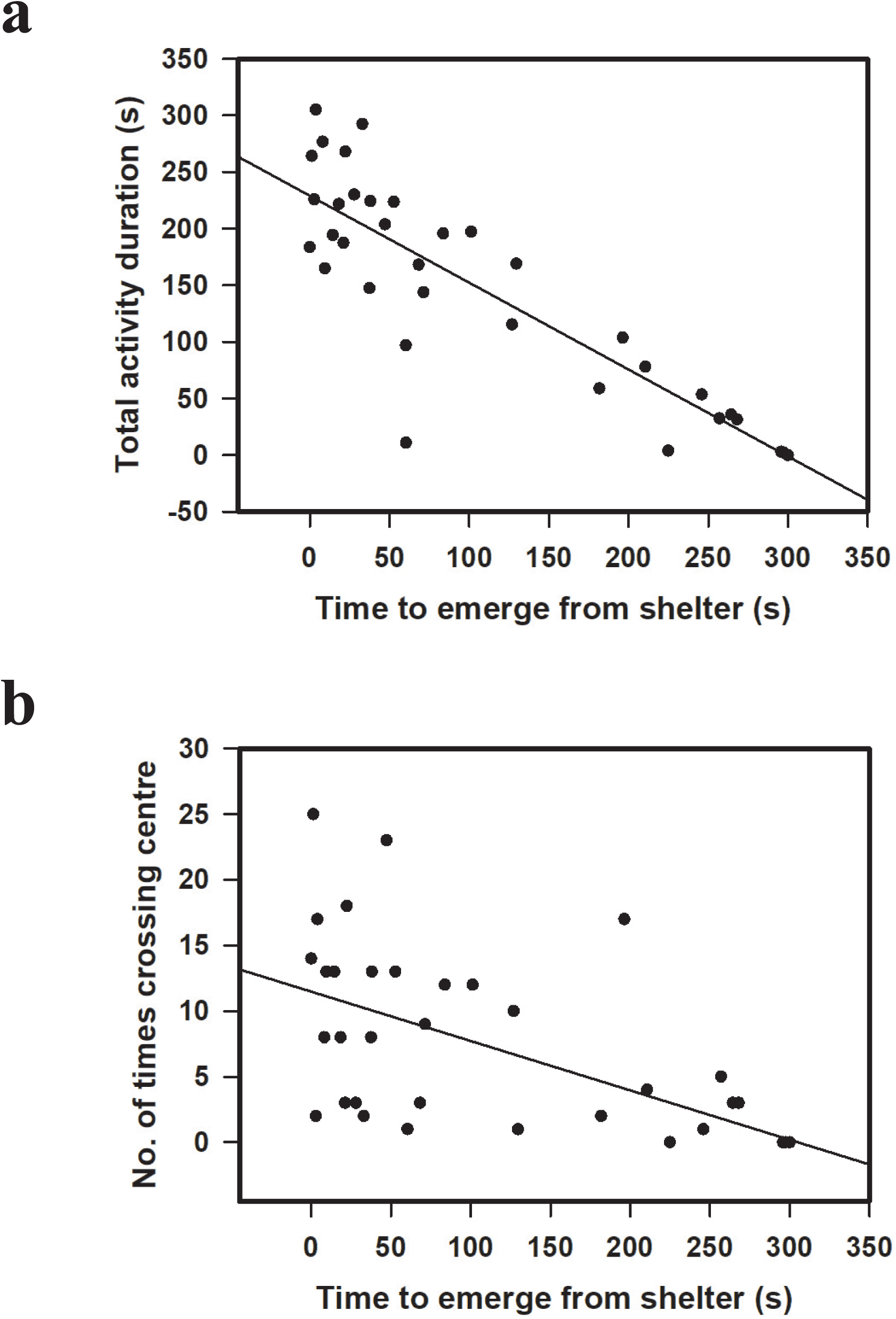
Correlations between boldness and activity in the juveniles of Deccan mahseer. a) Correlation between activity (in seconds) with latency to emerge from shelter (in seconds) which indicates boldness (bolder individuals emerge sooner; Spearman’s R = −0.96, p < 0.001). b) Correlation of number of times the fish crosses the centre (frequency) with latency to emerge from shelter (Spearman’s R = −0.88, p < 0.001). Each dot represents individual fish (n = 62).

**Figure 2.**
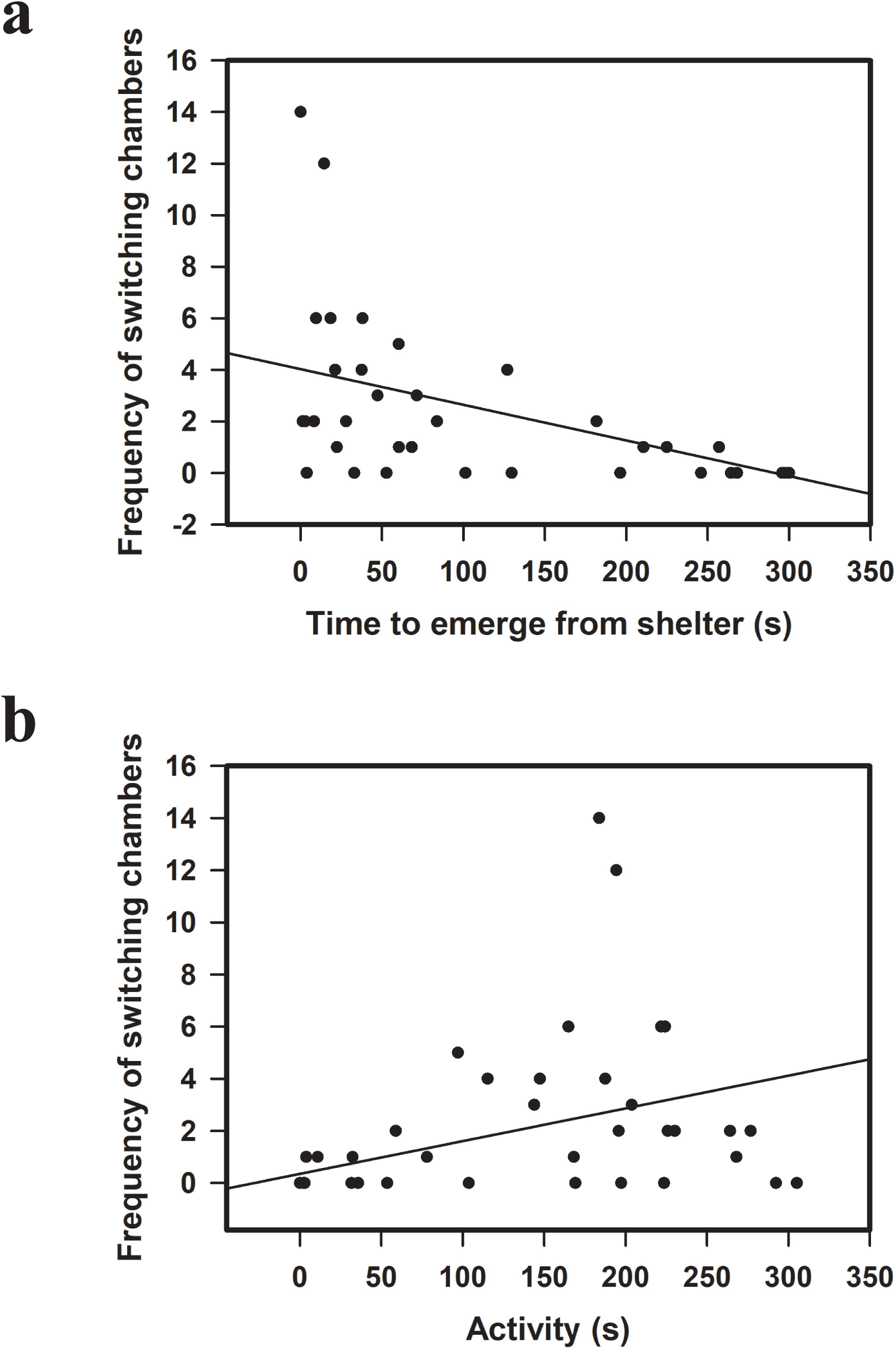
Correlations between switching chambers and other personality traits in the juveniles of Deccan mahseer. a) Correlation between number of times the fish switches between the two chambers and latency to emerge from shelter (Spearman’s R = −0.77, p < 0.001). b) Correlation between number of times the fish switches between the two chambers and total swimming activity (Spearman’s R = 0.66, p < 0.001).

### Sociability and lateralized utilization of brain hemispheres

Cross-context comparison of sociability revealed a significant effect of treatment (□^2^ = 18.96; p < 0.001; Kruskal-Wallis test). *Post hoc* analysis confirmed that sociability (time spent in proximity) exhibited towards conspecifics was significantly greater than the time spent near heterospecifics (Dunn test; p < 0.01) or empty chamber (p < 0.001; Figure 3a). Similarly, sociability towards heterospecifics was greater than sociability towards empty chamber (p < 0.05). No significant preference for left or right eye was noticed while the subject fish observed conspecific (p > 0.05; One-sample Wilcoxon signed rank test against 0), heterospecific (p > 0.05; Figure 4a) or the empty stimulus cage (p > 0.05). Cross-context comparison of LI using Kruskal-Wallis revealed no significant effect of treatment (□^2^ = 0.26; p > 0.05) suggesting no difference in lateralization in viewing empty stimulus cage, conspecifics or heterospecifics.

**Figure 3.**
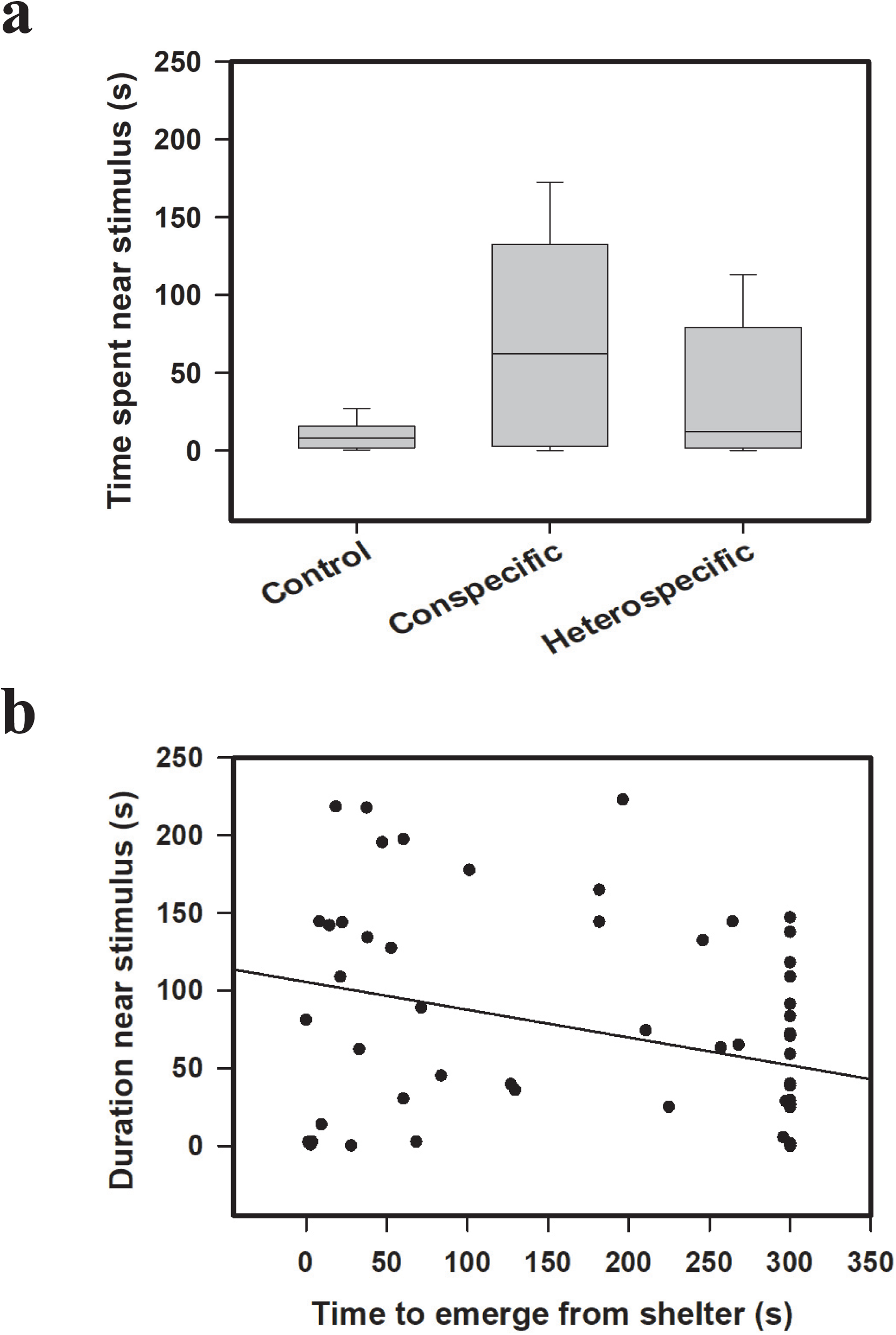
Sociability and its correlation with boldness in the juveniles of Deccan mahseer. a) Total duration of time (in seconds) spent by focal fish within 5 cm of the stimulus cage when cage was empty (Control), had a fish of the same species (Conspecific) or of a different species (Heterospecific). b) Correlation between sociability of fish towards conspecific measured by duration of time spent near stimulus cage and time to emerge from shelter (boldness; Spearman’s R = −0.34, p < 0.01).

**Figure 4.**
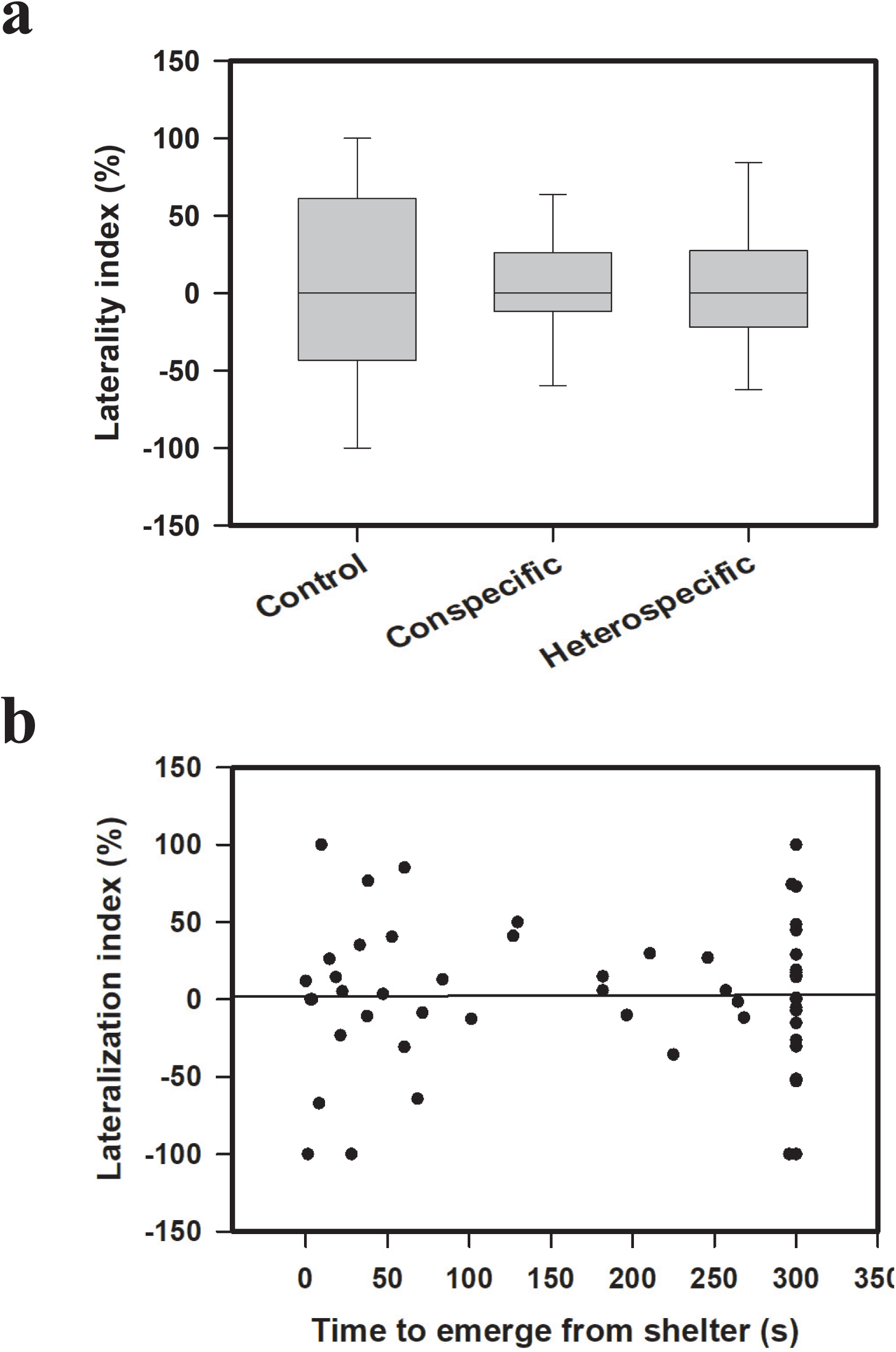
Lateralization and its correlation with boldness in the juveniles of Deccan mahseer. a) Lateralization index (in %) which is a measure of the preference of left eye over right eye (calculated as described in methods) to view the stimulus when focal fish is within 5 cm of the stimulus cage when cage was empty (Control), had a fish of the same species (Conspecific) or of a different species (Heterospecific). b) Correlation of laterality of fish when interacting with a conspecific measured by lateralization index with time to emerge from shelter (Spearman’s R = −0.008, p > 0.05).

Linear mixed models on sociability (log transformed) data as response variable revealed a significant effect of boldness (β = −0.003; p < 0.05), activity (β = −0.005; p < 0.05) and the presence of the conspecific (β = 0.67; p < 0.001). This suggests that emergence latency (boldness) and activity are negatively correlated with sociability (Figure 3b) while fish exhibited higher sociability towards conspecifics than empty chamber. The effect of heterospecific tilapia was not significant (β = −0.24; p > 0.05) suggesting that subject fish do not exhibit greater sociability towards Tilapia compared to the empty chamber. However, the interaction between the time taken to come out of the start chamber (boldness) and the time spent near heterospecific (β = 0.002; p < 0.01) was statistically significant with a positive coefficient. Thus, the positive correlation between sociability and boldness observed when fish is with a conspecific, is reversed by the presence of the heterospecific. This suggests that bold individuals are also more likely to be sociable towards their own species but not towards the other species. Linear mixed models of laterality did not reveal significant effects of boldness (β = −0.15; p > 0.05) and activity (β = −0.17; p > 0.05) or the identity of the stimulus fish (conspecifics: β = −9.4; p > 0.05; heterospecifics: β = −26.2; p > 0.05) suggesting that variation in lateralization is not associated with the personality of the individual in this species.

## Discussion

The relationship between big five animal personality traits has been widely studied across populations of different piscine species (Huntingford, 1976; Budaev, 1997; Burns, 2008). Although popular model systems such as sticklebacks, guppies and European wrasse exhibit strong correlation between boldness and exploratory activity in novel environments, later studies pointed to the role of ecological factors - the exposure to predators, resource availability and state-dependent safety (Bell, 2005; Dingemanse et al., 2007; Luttbeg and Sih, 2010) in determining the relationship between these traits. Interestingly, in nine-spined sticklebacks such an association was not seen (Herczeg et al., 2009) while brown trouts exhibited it only at certain life stages (Adriaenssens and Johnsson, 2013). The present study revealed that bolder juveniles of Deccan Mahseer would emerge into the swim-way earlier, tend to show greater activity and explore the well-lighted central areas of the apparatus more frequently than their shy counterparts; a result consistent with studies on several fishes across families (Conrad et al., 2011). In this context it should be noted that the individuals used in the present study lived in homogenous environment throughout their life and had no exposure to the predators which indicates the robust nature of the boldness-activity relationship in this species. However, contrary to the notion that the bold fishes will exhibit less time and reduced frequency of visiting start chamber, which may be functioning as a refuge area (Toms et al., 2010), bolder mahseers were found switching between chambers more often. This divergence suggests that more bold and active individuals explore all areas available in the swim way which is thus, expressed as higher frequency of visiting the start chamber. A similar response of greater chamber switching by bold individuals had been reported in gilthead sunbream *Sparus aurata* (Herrera et al., 2014) and round gobies (Marentette et al., 2011; 2012).

In many fish species, boldness exhibits a negative relationship with another axis of animal personality –sociability (Cote et al., 2010). Bold individuals show lower tendency for shoaling with others, often leave their group and extend their activities to novel areas of their habitat which could help them to find new sources of biologically significant materials at the cost of enhanced predation pressure. Meanwhile shy individuals prefer to remain with their shoal mates since large group size helps them to reduce the predation risk via dilution effect, early detection of the predators, etc. (Magurran and Pitcher, 1987; Lima and Dill, 1990; Hager and Helfman, 1991). In juveniles of Deccan mahseer, boldness had a positive relationship with sociability similar to that reported in feral populations of guppies (Irving and Brown, 2013) and bold individuals spent more time near conspecific in comparison to their shy counterparts. According to Irving and Brown (2013) such an association between boldness and sociability was restricted to one sex, males, in guppies. However, due to the lack of visible sexual dimorphism and the maturity it was difficult to identify the gender of the subject juvenile mahseers used in the present study. Although bold juvenile mahseers show increased activity in a novel environment, their greater sociability towards conspecifics can influence spreading of the populations from the site after release in to the natural habitats. According to Cote et al. (2010), it is the bold individuals which establish the new colonies after reaching a new ecosystem due to their enhanced tendency to take risk. Additionally, juvenile mahseers continuing as tight shoal in an unfamiliar habitat can attract the attention of the predators. However it should be noted that the individuals tested in this study are only 2 months old and the relationship between boldness and sociability may change once the individual grows. For instance, correlations between boldness and aggression seen at the adult stage in sticklebacks are weaker during the sub-adult stage (Bell and Stamps, 2004). Hence a continuous evaluation of the linkage between boldness and sociability in different stages of growth is essential to find the suitable age at which this fish could be released into the natural habitats to establish viable populations with maximum survival.

Young ones of many social piscine species are known to show high shoaling tendency due to the heightened vulnerability for predation and some fish even form part of multi-species groups for the same reason (Landeau and Terborgh, 1986; Overholtzer and Motta, 2000). However, living with exotic invader species can have undesirable effects on the behaviour and personality traits of the indigenous species (Blanchet et al. 2007). Moreover, invasive species can acquire information from native species and effectively invade habitats while shoaling with them (Sievers et al., 2012; Camacho-Cervantes et al., 2015). The changes induced by tilapia on the boldness-sociability relationship of the juvenile mahseers and spending significantly more time close to heterospecific fish over the empty presentation cage raises many ecological concerns. The highly sociable mahseers may try to join heterospecific shoals in the absence of conspecifics and individuals getting isolated after release into natural water bodies during the time of predator attack or monsoon flood may be unavoidable. Becoming part of a shoal with heterospecifics could result in a substantial cost of competition for food and also increase detection by predators due to the oddity effect (Ward et al., 2002). Furthermore, increasing presence of many alien invasive species has been reported from the natural habitats of Decan mahseers (Kharat et al., 2003) and there is still little information on how the individuals varying in their personality traits adapt with the pressure exerted by such exotic species. Although juveniles of Deccan mahseer exhibited significant difference in their sociability towards conspecific and exotic heterospecific, no such variation in the lateralized utilization of brain hemispheres while observing another mahseer individual or tilapia was noticed. The absence of, or reduction in lateralisation is often associated with cognitive deficiencies and impairment of synchrony in shoaling behaviour and hence could impact the fitness of the individuals in natural habitats (Bisazza and Dadda, 2005; Dadda et al., 2010; Bibost and Brown, 2014; Frasnelli and Vallortigara, 2018). Although the current study revealed a lack of significant asymmetry in the brain use by juvenile Deccan mahseers while observing a social stimulus, it is not clear whether it is a characteristic feature of this species or the result of living in a homogenous environment without any predation pressure. Reduced degree of lateralization has been noted in the populations of rainbowfish reared in conditions of minimal environmental complexity (Bibost et al., 2013) as well as in populations of *Brachyraphis episcopi* collected from areas with lower predation pressure (Brown et al., 2004). Additionally, lack of significant correlations between laterality, sociability, activity and boldness could also be due to measuring lateralization in an unfamiliar arena since previous studies on convict cichlids and Port Jackson sharks have shown link between laterality and boldness in familiar but not unfamiliar environment (Reddon and Hurd, 2009; Byrnes et al., 2016). The expression of lateralization is also more pronounced in a familiar environment perhaps because the fish are still assessing the environment when first exposed to novel conditions before establishing preferred directionality (Byrnes et al., 2016). These results also emphasize the need for testing individual and population level lateralization in Deccan mahseer in contrasting environmental conditions in the laboratory and in natural habitats to get a better picture of personality-laterality relationship in this species.

## Conclusion

The results of the current study exploring individual variation in major personality traits, lateralized utilization of brain hemispheres and relationship between personality and laterality have wider implications for the *in situ* and *ex situ* conservation and cultivation of the endangered mega fish Deccan Mahseer. Although thousands of juveniles of this species reared in hatchery are released into the natural habitats to replenish the stock, selection of individuals with personality traits most suitable for the target ecosystem could be useful in enhancing their survival (Reale et al., 2010; Conrad et al., 2011). According to Ullah et al. (2017) young ones of another species of Mahseer, the Himalayan Mahseer (*Tor putitora*) exhibited enhanced boldness, exploratory behaviour, antipredator response etc. when reared in spatially enriched environments. Unfortunately, such lines of research focusing on Deccan mahseer as subject species are yet to commence. Additionally, tracking patterns of interaction of individual mahseers divergent in their personality traits towards alien invasive piscine species in isolation and in shoals is also an urgent requirement since the natural habitats of mahseers are getting invaded with such species. Hence it is expected that future studies on Deccan Mahseer will elucidate a clear picture of the behavioural and cognitive abilities of this species and contribute to the efforts to save this species from extinction.

## Acknowledgements

The authors would like to thank Director, Karnataka Fisheries Department for kindly providing the fingerlings of Deccan Mahseer for research purposes. We are grateful to Darshan Pramod, District Fisheries Department Kodagu for her support. Financial support from the DBT-RA Program in Biotechnology and Life Sciences to Vishwanath Varma is gratefully acknowledged.

